# Conformational diversity of CDR region during affinity maturation determines the affinity and stability of Sars-Cov-1 VHH-72 nanobody

**DOI:** 10.1101/2020.12.08.416164

**Authors:** Suji George

**Affiliations:** Independent Researcher

**Keywords:** antibody, nanobody, affinity maturation, somatic hypermutation, stability, flexibility, molecular dynamics

## Abstract

The affinity maturation of Sars-Cov-1 VHH-72 nanobody from its germline predecessor has been studied at the molecular level. The effect of somatic mutations accumulated during affinity maturation process on flexibility, stability and affinity of the germline and affinity matured nanobody was studied. Affinity maturation results in loss of local flexibility in CDR of H3 and this resulted in a gain of affinity towards the antigen. Further affinity maturation was found to destabilize the nanobody. Mechanistically the loss of flexibility of the CDR H3 is due to the redistribution of hydrogen bond network due to somatic mutation A50T, also this contributes significantly to the destability of the nanobody. Unlike antibody, in nanobody the framework region is highly conserved and structural diversity in CDR is the determining factor in diverse antigen binding and also a factor contributing to the stability. This study provide insights into the interrelationship between flexibility, stability and affinity during affinity maturation in a nanobody.

## Introduction

Antibodies display exquisite molecular recognition properties and are the effector molecules of the humoral immune system. They are widely used in biological research and also find application in biopharmaceutical industry due to its high specificity towards target antigens. The number of potential foreign antigen a person could be exposed in a lifetime outnumbers the germline repertoire formed by the V, D and J gene recombination. Using 69 VH, 27DH and 6 JH, gene segments, the immune system generates 11,000 unique VDJ gene segments. The antibody repertoire of an individual is estimated to be populated with 10^11^ antibodies formed by recombination events of the above mentioned heavy chain genes and 31 IGKV genes, 5 IGKJ genes, 45 IGLV genes and 7 IGLJ genes that comprise light chains (Cisneros et al., 2019)□. Besides recombination event, affinity maturation by somatic mutation is another mechanism of diversifying the specificity towards the antigen.

Other formats of antibodies such as the single chain fragment variable, Fab and nanobodies (Nbs) are potential tractable class of antibody derived fragments having application in biological research, biopharmaceuticals and diagnostics. Nanobodies are class of heavy chain antibodies found in camlid species such as camels, llamas and alpacas lacking the light chain. They are almost ten times smaller than conventional antibody with molecular weight ~15 kDA and retains comparable binding specificity (Muyldermans, 2013). Further due to its small size Nb can access hidden epitopes, which otherwise is inaccessible to conventional antibodies in enzyme active sites, virus capsid and G-protein coupled receptors. Similar to the Variable heavy chain (VH) domain of the antibody the VHH domain of the Nbs are composed of four framework regions (FR1, FR2, FR3, FR4) and three complimentary determining region (H1, H2, H3). Inspite of the absence of variable light chain (VL), Nb shows remarkable similarity in specificity like the counterpart antibody (Muyldermans, 2013)□. In antibody the paratope lies at the interface between the VH-VL and regions from all the six hyper variable loop regions contributes to its binding to the antigen. In addition the VH-VL orientation also plays a crucial role in determining the shape of the antigen-antibody interface, while in Nbs the paratope is contained within the VHH domain with considerably lower antigen-antibody surface. The remarkable binding specificity in Nbs is thus achieved by its small size and only single domain architecture.

A comparison of the FR in nb and antibody shows that frame work region is highly conserved both in sequence and structure than Abs suggesting that it is not widely used for generation of diverse binding specificities (Mitchell and Colwell, 2018)□. Also unlike the antibodies, the residues corresponding to the interface in Nbs shows less sequence variation and is therefore suggested to be not under evolutionary pressure. Thus a plausible mechanism by which Nbs generates diverse range of binding specificity is through the sequence and structural diversity of its CDR loops. Another mechanism by which Nbs compensates for the absence of VL chain is by drawing the paratope residues from a larger set than antibodies there by favoring diversity in shape and physical properties of the paratope region (Mitchell and Colwell, 2018)□. The VHH gene is derived from the IGHVH family of V genes which is hypothesized to be evolved via the duplication of a single IGHV3 gene family. However in case of antibody heavy chains have been identified with IGHV4 and IGHV3 V genes with diverse pool of starting germline sequence. Thus if VHH is derived from such a considerably small pool of germline sequence, then there is possibility that the starting germline is less variable in the region encoded by the V gene namely the FR 1 FR2 FR3 H1 and H2 (Muyldermans, 2013). Thus affinity maturation in Nbs becomes more important than in antibodies to generate diverse range of binding specificity through structural variation.

Studies suggest that germline antibodies from being conformationally diverse progress towards more specific and conformationally restricted variants by affinity maturation (Ovchinnikov et al., 2018; Wedemayer, 1997). In the absence of VL in Nbs and with limited sequence and structural diversity in FR, the loop region plays crucial role in generating diverse binding specificities. It is thus the objective of the study to probe the role of affinity maturation in the conformational diversity of Nbs. For this we compared the molecular dynamics of germline and mature nanobody. Further using systematic stability analysis of specific single point mutant Nb it has been show that the mutation in loop region is indeed important than the FR region in determining the stability and affinity.

## Methods

### Nanobody Modeling

The X-ray crystal structure of the Sars-Cov-1 Receptor Binding Domain (RBD) bound by the single domain antibody SARS VHH-72 antibody (6waq was downloaded from the protein Data Bank (PDB; www.pdb.org (Wrapp et al., 2020)□. The hetero atoms were removed and residues in the PDB were sequentially renumbered. For modeling the germline counterpart, sequences were retrieved by homology search using V-Quest at IMGT (Brochet et al., 2008)□. The amino acid of germline heavy chain V, D and J gene fragments were then manually assembled to obtain the full length nanobody germline sequence. Germline Nb were modeled using the AbodyBuilder server (Leem et al., 2016). AbodyBuilder selects the templates using the sequence identity of the framework regions and the program modeler then models the FR regions. The CDR regions are modeled using the Fread, loop prediction algorithm by searching against a CDR database. The query sequence is searched against the SAbDab (Structural Antibody Database) (Dunbar et al., 2014)□, which also has VHH and modified VH/VL existing as single domain antibodies. The PDB code of the templates for 6waqG was B chain of the structure 6imk and throughout the manuscript germline Nb structure would be designated by 6waqG and mature Nb structure with 6waqM.

### Molecular dynamics

MD calculations were done using the program gromacs-2019.4 using the amber99sb-ildn force field available in gromacs (Van Der Spoel et al., 2005). The nanobody was explicitly solvated in a cubic box with a distance of 1.2 nm from the structure to the edge of the box, using the periodic boundary condition (pbc= xyz) and using the water model TIP3P. Further the system was neutralized by adding sodium and chloride ions. The structure was then subjected to constrained energy minimization using the steepest descent algorithm with step size of 0.01 nm along the gradient and maximum of 10000 steps. All short ranged non bonded interactions were truncated at 1.0 nm (rcoulomb=1.0 and rvdw=1.0). The particle mesh ewald method was used to calculate the long range electrostatic forces. Following energy minimization the structure was equilibrated in two steps process. During the first step 100 ns NVT ensemble equilibration with applied position restraints the system was coupled to thermostat with reference temperature of 300K using the velocity rescale scheme (Bussi et al., 2007). This was followed by 500 ns NPT ensemble equilibration step with position restrain maintained on the protein non hydrogen atoms with reference pressure 1 bar using the barostat method proposed by Parrinelo and Rahman (Parrinello and Rahman, 1981)□. After the equilibration the MD simulation was extended to 100ns without positional restraints at 300K. Similar step as mentioned above was carried for simulation of the nb at 500K for partial unfolding studies. Adequate quality control measure during the energy minimization and equilibration stage of MD analysis was adopted before finally starting the production run. After production run detailed analysis including RMSD, RMSF, radius of gyration, cluster analysis was carried out using the inbuilt tools in gromacs. Visualization of the trajectory and figures were made using the Chimera (Pettersen et al., 2004)□ and VMD (Humphrey et al., 1996)□.

### Stability calculation for Germline and Mature Nanobody

The effects of somatic hyper mutation on the stability of Nbs were studied using FoldX. The FoldX “RepairPDB” command was used to correct the structure of the Nbs (Delgado et al., 2019)□. Further to delineate the contribution at each position single point mutant model was build by “BuildModel” command. The following equation 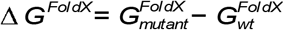 where 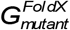 is the mutant FoldX score and 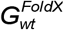 is the wild type FoldX score used to to obtain the change in stability.

### Docking of nanobody to RBD

We used the web interface of RosettaDOCK to dock 6waqG and 6waqM to RBD (Chaudhury et al., 2011; Lyskov and Gray, 2008)□□. The ΔG of binding and other interface parameters were calculated using the Rosetta set of programs (Alford et al., 2017)□. In addition we also used the prodigy web server for calculation of binding affinity of the complex (Kastritis and Bonvin, 2012).

## Results and discussion

### Modeling of germline nanobody

Antibody and antigen complex are ternary structures forming three-way interactions. The light chain (LC) of the antibody interacts with heavy chain (HC) and both the HC-LC chain complex interacts with the antigen. Thus the secondary interface in variable region of antibody formed by various orientations of HC and LC determines the geometry of the paratope during antigen recognition. Somatic hyper mutation in the VH-VL interface during affinity maturation can improve the binding affinity independent of the affinity maturation at the antibody antigen interface. Although not a universal phenomenon, affinity maturation in some antibodies results in rigidification of the variable region which has been suggested to improve binding characteristics of the antibody. Mutation at HC-LC interface induces alternate orientation through a combination of shifting the energy minimum and limiting the amount of conformational entropy by stabilising the HC-LC interface in a way that favors the bound conformation (Cisneros et al., 2019). Unlike antibody, nanobody is devoid of the HC-LC interface, since it has only the HC. Thus in the absence of LC, the study of the effect of somatic hypermutation of nanobody on its flexibility and interaction with antigen would add further advancement of the knowledge about antibody antigen interaction. For this study we chose single domain antibody SARS VHH-72, known to bind to the RBD region of the spike protein of Sars-Cov-1. For modeling of the germline counterpart sequences were retrieved by homology search using V-Quest at IMGT (Brochet et al., 2008; Giudicelli et al., 2011)□. The DNA sequences of the germline heavy chain the V, D and J gene fragments were manually assembled and then translated into the amino acid sequence. The nanobody is derived from the germline gene Vicpac IGHV-3-3*01 F (V region), Vicpac IGHJ4*01 F (J-region) and Vicpac IGHD5*01 F (D-region). The x-ray crystal structure 5imk B chain was used as template by the Abodybuilder to model the FR and CDR region of 6waqG.. Molprobity analysis a validation method for experimental or predicted protein models with metrics such as the clash score, poor rotamers, Ramachandran outliers, Ramachandran favored and a molprobity score (Chen et al., 2010) was used to validate the germline structure. The models were further evaluated by measuring the C_α_ backbone RMSD between the 6waqG model vs 6waqM (1.30 Ǻ) and 6waqM and template 5imk X-Ray (1.4 Ǻ) crystal structure. We did not find any difference in the RMSD between 6waq-G and 5imk as expected 5imk is the template for modeling 6waq-G. The region wise RMSD, the CDR H3 (18 residues) shows the highest RMSD of 2.8 followed by CDR H1 (13 residues) with RMSD 1.4 Ǻ followed by CDR H2 (10 residues with RMSD 0.85 Ǻ. Figure-1 shows the alignment of sequence of 6waqM with 6waqG and the mutation acquired during affinity maturation with mutation V5Q in FR-1, S30E in CDR H1, A50T in CDR H2, A61T E89D in FR-3 (Table 1). Table 2 shows the residue of 6waq-M in contact with RBD of the spike protein with 14 residues of CDR-H3, 10 residues of CDR-H2, 4 residues of FR-3 and 2 each of CDR1 and FR2.

**Figure 1:**
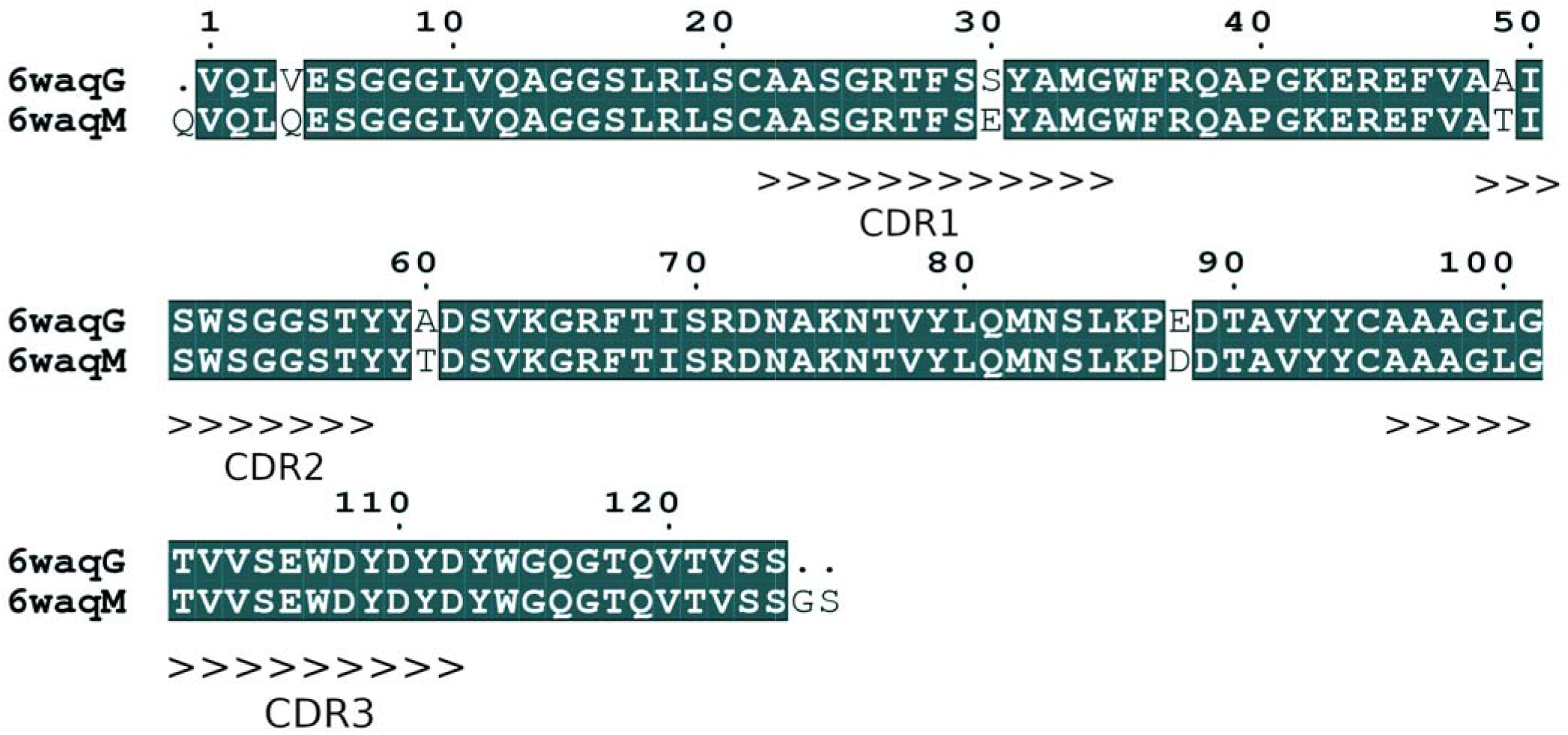
**A)** Alignment of 6waqG and 6waqM showing the sites of somatic mutation and CDRs

**Table 1:** Sites of somatic mutation in 6waqG and 6waqM nanobody. PDB was numbered by 1) IMGT scheme 2) Sequentially

|  | Mutation 6waqGVs 6waqM<br>(IMGT numbering) | Mutation 6waqG Vs 6waqM<br>(PDB numbering) |
| --- | --- | --- |
| FR1 | V5Q | V5Q |
| FR2 |  |  |
| FR3 | A68T E97D | A61T E89D |
| FR4 |  |  |
| H1 | S36E | S30E |
| H2 | A55T | A50T |
| H3 |  |  |

**Table 2:** Interface residues in 6waqG-RBD and 6waqM-RBD complex.

|  | **6waqM.pdb** | **6waqG.pdb** |
| --- | --- | --- |
| FR1 |  |  |
| FR2 | 46-E and 47-F | 39-Q, 40-A, 41-P, 42-G, 43-K,<br>44-E, 45-R, 47-F |
| FR3 | 60-Y, 61-T, 62-D, 65-K | 61-A, 62-D |
| FR4 |  |  |
| H1 | 31-E and 33-A |  |
| H2 | 50-T, 51-I, 52-S, 53-W, 54-S, 55-G, 56-G, 57-S,<br>58-T, 59-Y |  |
| H3 | 99-A, 100-G, 101-L, 102-G, 103-T, 104-V, 105-V,<br>106-S, 107-E, 108-W, 109-D, 110-Y, 111-D,<br>112-Y | 104-V, 105-V, 107-E, 108-W,<br>109-D, 110-Y, 111-D, 112-Y |

### Flexibility changes in nanobody

Root mean square fluctuation (RMSF) is a widely used metric to measure on average fluctuation in dynamics of a residue within a structure. The RMSF comparison between 6waqG and 6waqM antibody reveals that the effect of somatic hypermutation is local confined to the H3 and region of S30E mutation in H1 region (Figure 2A). Further it shows that in 6waqM there is considerably lower fluctuation in the H3 region and higher fluctuation at region around S30E mutation in H1 region. Thus the results that we present here is consistent with earlier studies of antibodies and with the traditional paradigm of affinity maturation, which shows a loss of flexibility during maturation in our case there is significant loss of flexibility in the H3 region (Ovchinnikov et al., 2018). In order to find out whether the protein secondary structure is stable during simulation at 300 K the average contact map of the protein was calculated, which clearly shows that the protein’s secondary structure is stable, with antiparallel packing between strands β1-β3, β3-β8, β4-β9, β4-β5, β5-β6, β7-β8, β9-β11, β9-β10, β4-β9, β4-β5, β5-β6 the parallel packing between strands β2-β11 maintaining contacts during 50ns simulation (Data not shown) (Mercadante et al., 2018). Further the data in figure 2B, 2C shows the standard deviation of the symmetric contact map with regions of high fluctuation and higher standard deviation of the inter-residue contact distance further confirmed the higher flexibility of H3 region in 6waqG.

**Figure 2.**
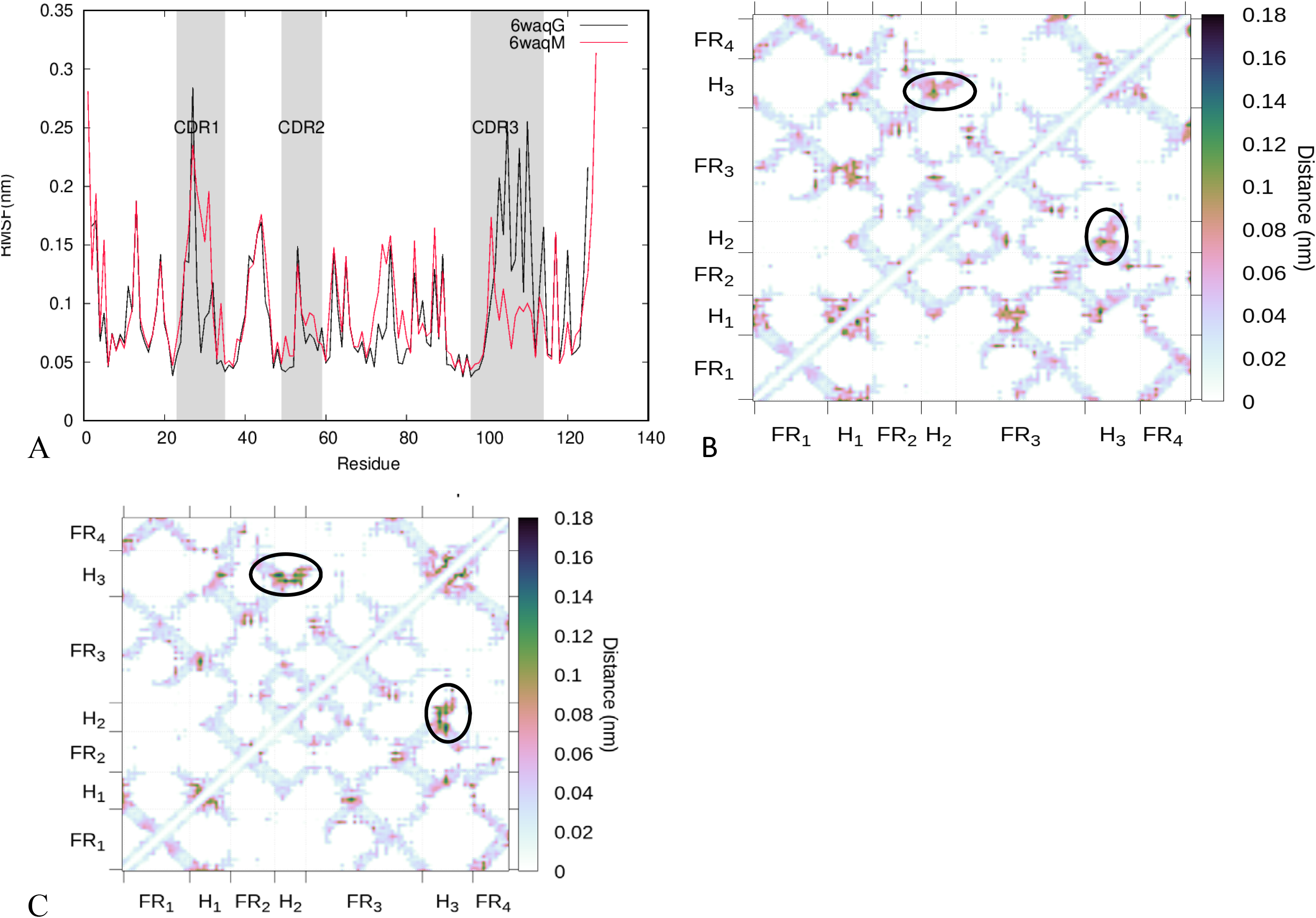
A) RMSF plot of 6waqG and 6waqM B) Symmetric contact map showing the standard deviation of contact distances along the 50 ns trajectory at 300K for 6waqG C) Symmetric contact map showing the standard deviation of contact distances along the 50ns trajectory at 300K for 6waqM.

### Stability of nanobody

The pearson correlation coefficient for inter residue distance for both 6waqG and 6waqM nbs was compared. Correlation of inter residue distances with time are shown with negative pearson coefficients (orange) correspond to contact formation and positive ones (purple) correspond to contact ruptures. The higher positive pearson coefficient in case of 6waqM compared to 6waqG indicates significantly more contact ruptures and to further substantiate this finding we performed stability analysis of 6waqM and single and double point mutants derived from 6waqG to find out the overall stability. Figure 3 shows that the affinity maturation overall leads to decrease in stability of the nanobody and the mutation at A50T contributes maximum to decrease in stability. To further find out region of lower stability we performed molecular dynamics at 500K to study the partial unfolding of the nanobody. Figure 4 shows the average local interaction time of β strands of the nbs. The region corresponding to β1, β9, β10 of 6waqM showed considerably lower average interaction time indicating region of lower stability. Thus the rigidification of the CDR3 region in 6waqM destabilizes β9, β10 strands.

**Figure 3:**
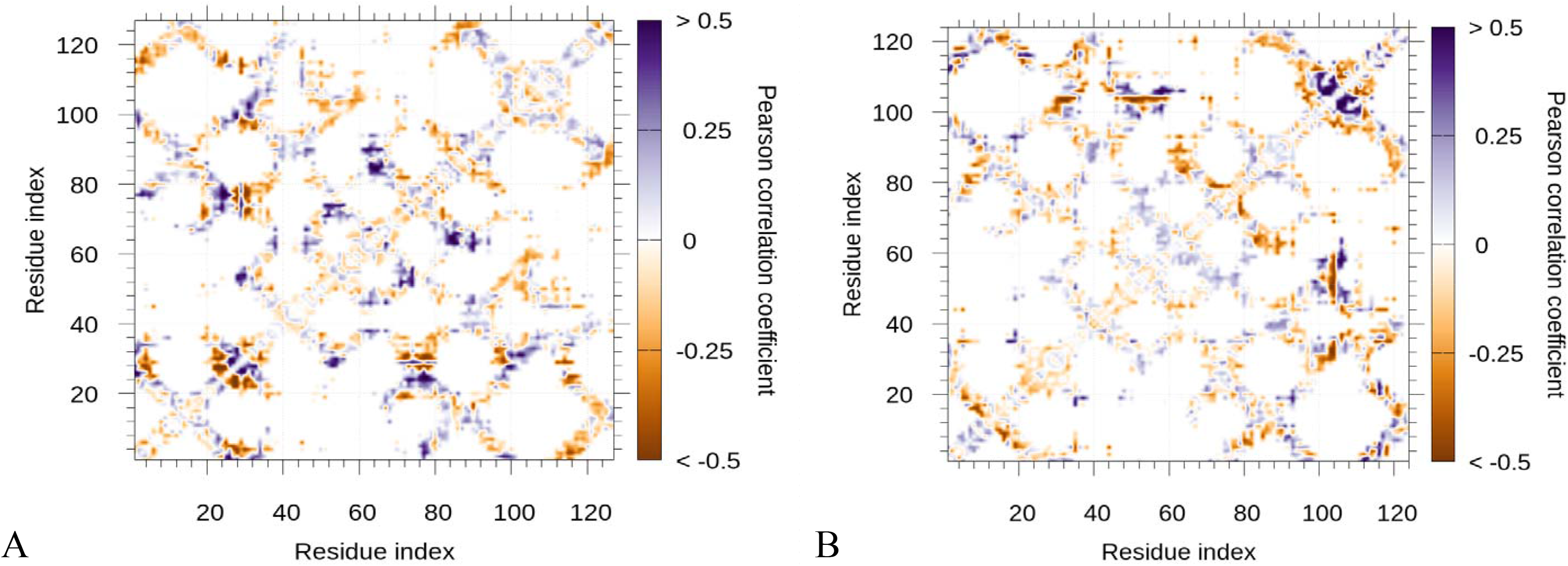
Correlation of interresidue distance with time. Negative pearson coefficients (orange) correspond to contact formation and positive ones (purple) correspond to contact ruptures. A) 6waqM B) 6waqG

**Figure 4:**
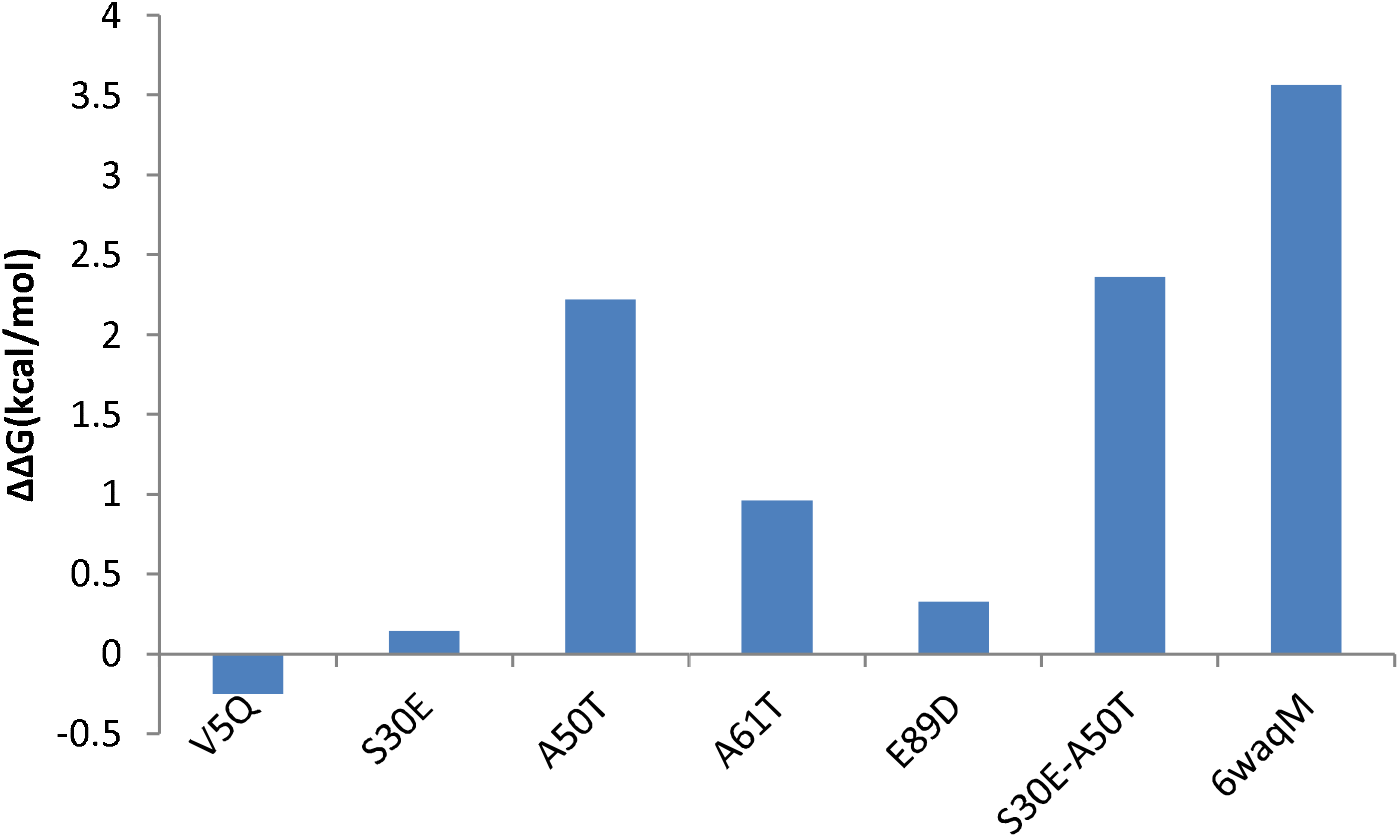
FoldX ΔΔG score for single, double point mutants and mature nanobody

### H-Bond network differences

In order to explain the differences observed in rigidity of 6waqG and 6waqM nbs we analysed the hydrogen bond network differences. Hydrogen bond (HBN) plays an important role in protein structure stability, dynamics and functions because they stabilize secondary structure and also plays a crucial role in forming the tertiary structure. As HBN is involved in constraining topology and there by fluctuation, probing the change in HBN network due to mutation can provide crucial clue regarding the differences in rigidity. For this we calculated the evolution of HBN during the trajectory and only the occupancy above 50% were considered for analysis. Table-3 shows the hydrogen bond acceptor and donor and corresponding occupancy. The largest hydrogen bond differences involving side chain between the germline and mature would explain the rigidity differences and a visual inspection reveals that the hydrogen bond between SER106-side ASP111-side with occupancy 81.58% and TYR-59 side in CDR H2 and GLU-107 side in CDR H3 with occupancy 59.6 % is present in 6waqM. The local HBN network around distance of 10 Ǻ of A50 in 6waqG and T50 in 6waqM is shown in figure 6. It can be seen that A50T somatic mutation in 6waqM alters the HBN network thereby constraining the CDR H3 region.

**Table 3:**
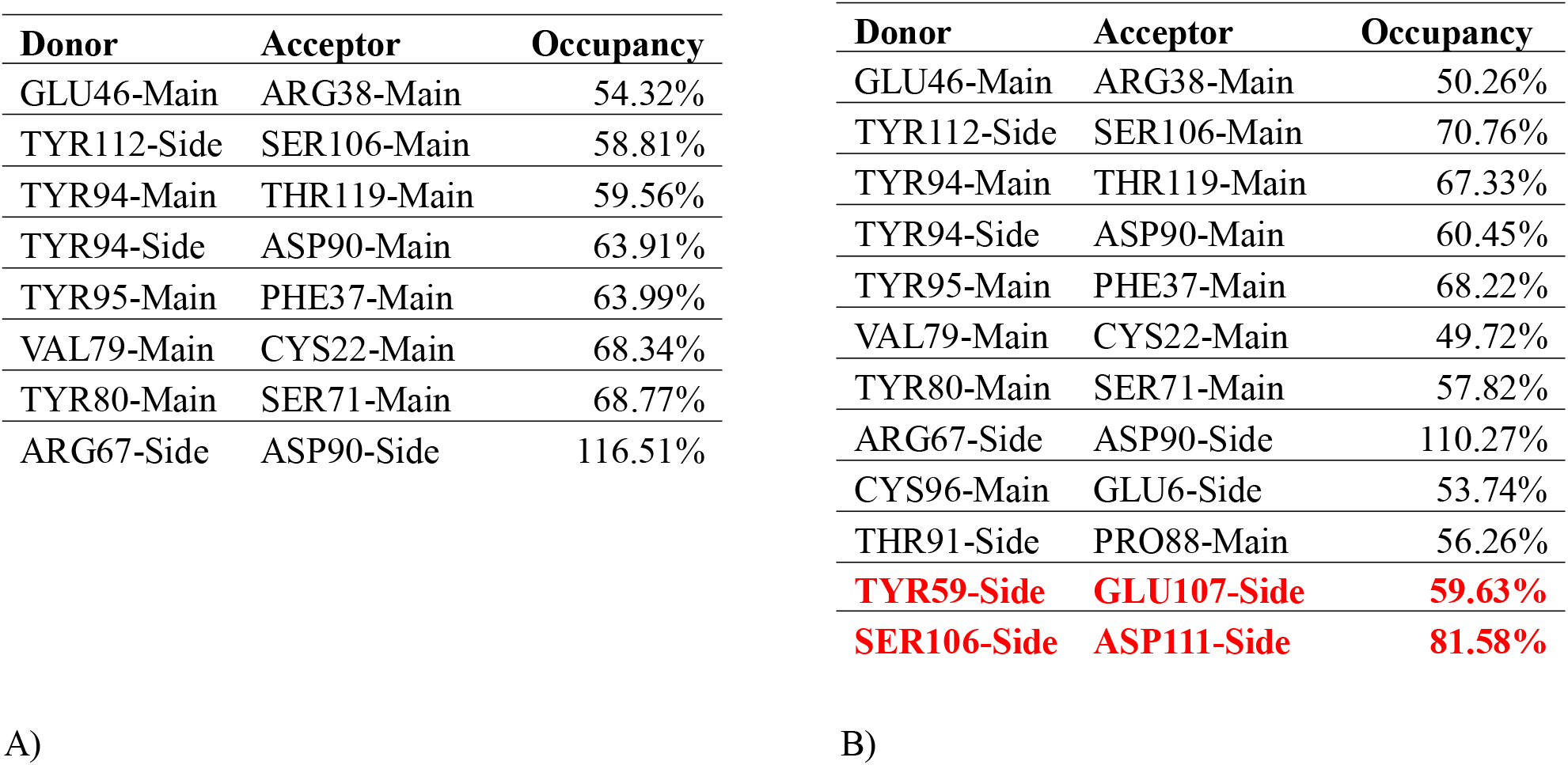
Intra-molecular H-bond donor, acceptor and occupancy A) 6waqG B) 6waqM

Table-2
| Donor | Acceptor | Occupancy |
| --- | --- | --- |
| GLU46-Main | ARG38-Main | 54.32% |
| TYR112-Side | SER106-Main | 58.81% |
| TYR94-Main | THR119-Main | 59.56% |
| TYR94-Side | ASP90-Main | 63.91% |
| TYR95-Main | PHE37-Main | 63.99% |
| VAL79-Main | CYS22-Main | 68.34% |
| TYR80-Main | SER71-Main | 68.77% |
| ARG67-Side | ASP90-Side | 116.51% |

A) Table-3
| Donor | Acceptor | Occupancy |
| --- | --- | --- |
| GLU46-Main | ARG38-Main | 50.26% |
| TYR112-Side | SER106-Main | 70.76% |
| TYR94-Main | THR119-Main | 67.33% |
| TYR94-Side | ASP90-Main | 60.45% |
| TYR95-Main | PHE37-Main | 68.22% |
| VAL79-Main | CYS22-Main | 49.72% |
| TYR80-Main | SER71-Main | 57.82% |
| ARG67-Side | ASP90-Side | 110.27% |
| CYS96-Main | GLU6-Side | 53.74% |
| THR91-Side | PRO88-Main | 56.26% |
| **TYR59-Side** | **GLU107-Side** | **59.63%** |
| **SER106-Side** | **ASP111-Side** | **81.58%** |

### Effect of rigidity and stability changes on antigen affinity

In order to find out whether loss of flexibility in CDR H3 region has any effect on the antigen binding ability docking was performed followed by binding affinity calculations. The results indicate that there is a gain in antigen binding affinity upon loss of flexibility with ΔG of binding −8.9 kcal/mol (Kd=3.00E-07 M) for 6waqM and −8.3 kcal/mol (Kd=7.70E-07M) (Figure-5). Consensus results as above were obtained using the rosetta interface analyzer and other interface metric such as the shape complimentarity of the interface Sc=0.619 for 6waqM and Sc=0.630 did not show significant difference. However the buried solvent accessible surface area (dSASA) shows significant difference between 6waqM and 6waqG. Table 2 shows the interacting residues at the interface with residues from FR2, FR3, CDR1, CDR2 and CDR 3 interacting with the RBD in 6waqM, while in case of 6waqG only the residues from FR2 and CDR3 interacts with RBD

**Figure 5:**
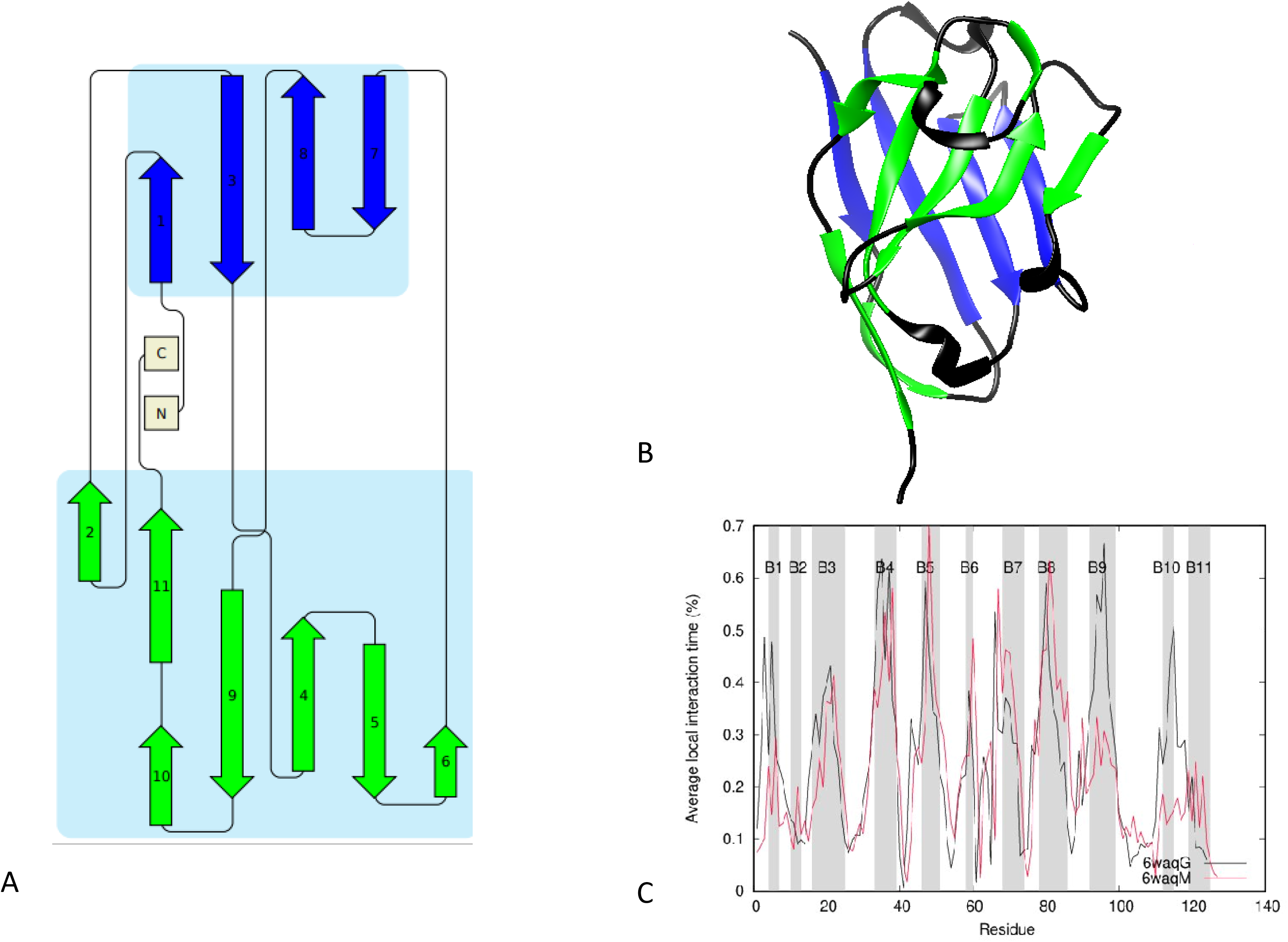
A) Schematic diagram of the mature antibody structure B) cartoon diagram of the 6waqM C) Average local interaction profile for 6waqG for simulation at 500K. B1, B2, B3 etc represents β1, β2, β3 strands.

**Figure 6:**
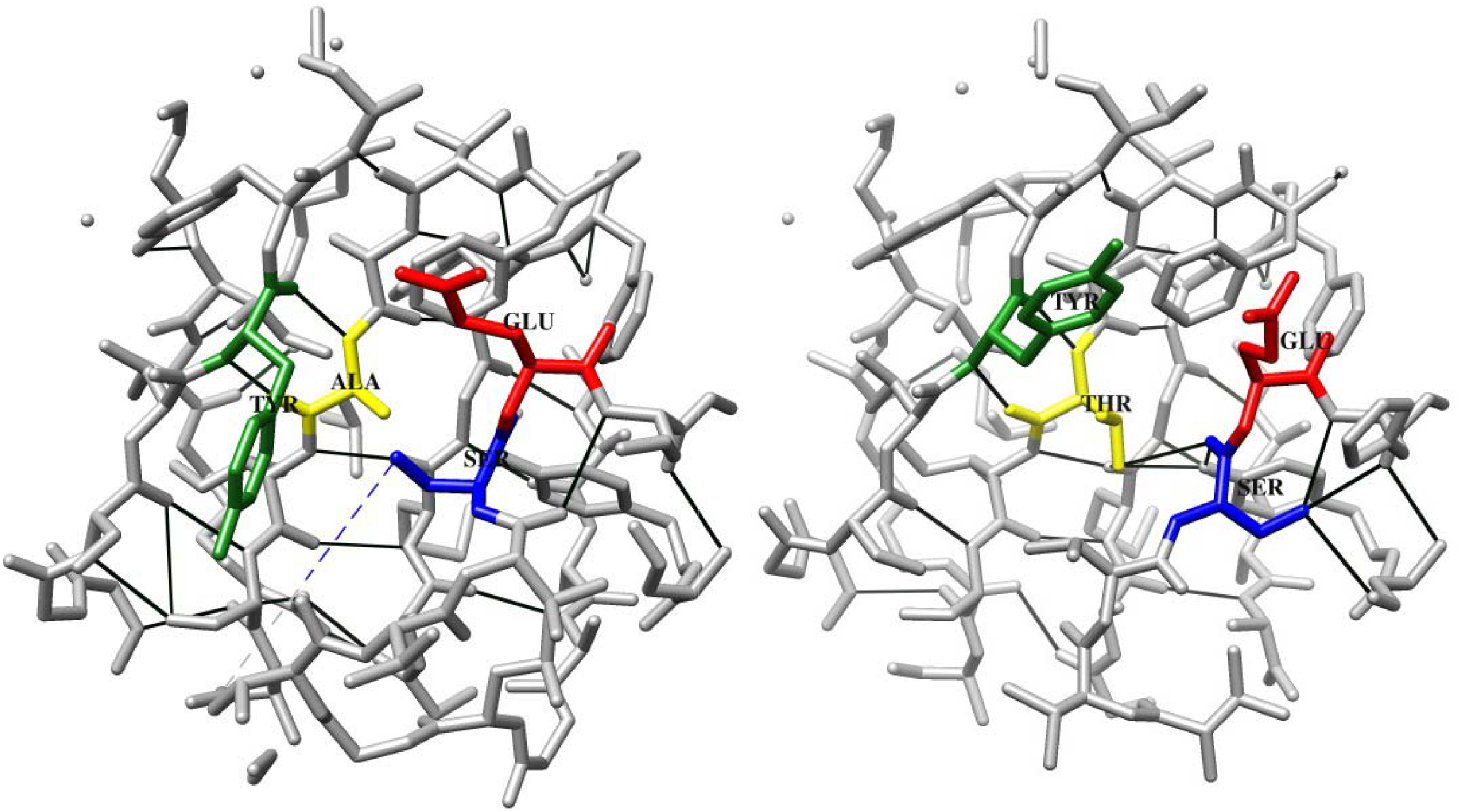
H-Bond network in 6waqG and 6waqM residues with in a distance of 10 Ǻ around the mutation site A50T site is shown. **Colour code**: Yellow-Alanine, Red-Glutamate, Green-Tyrosine Blue-Serine, Black-Hydrogen Bond

**Figure 7:**
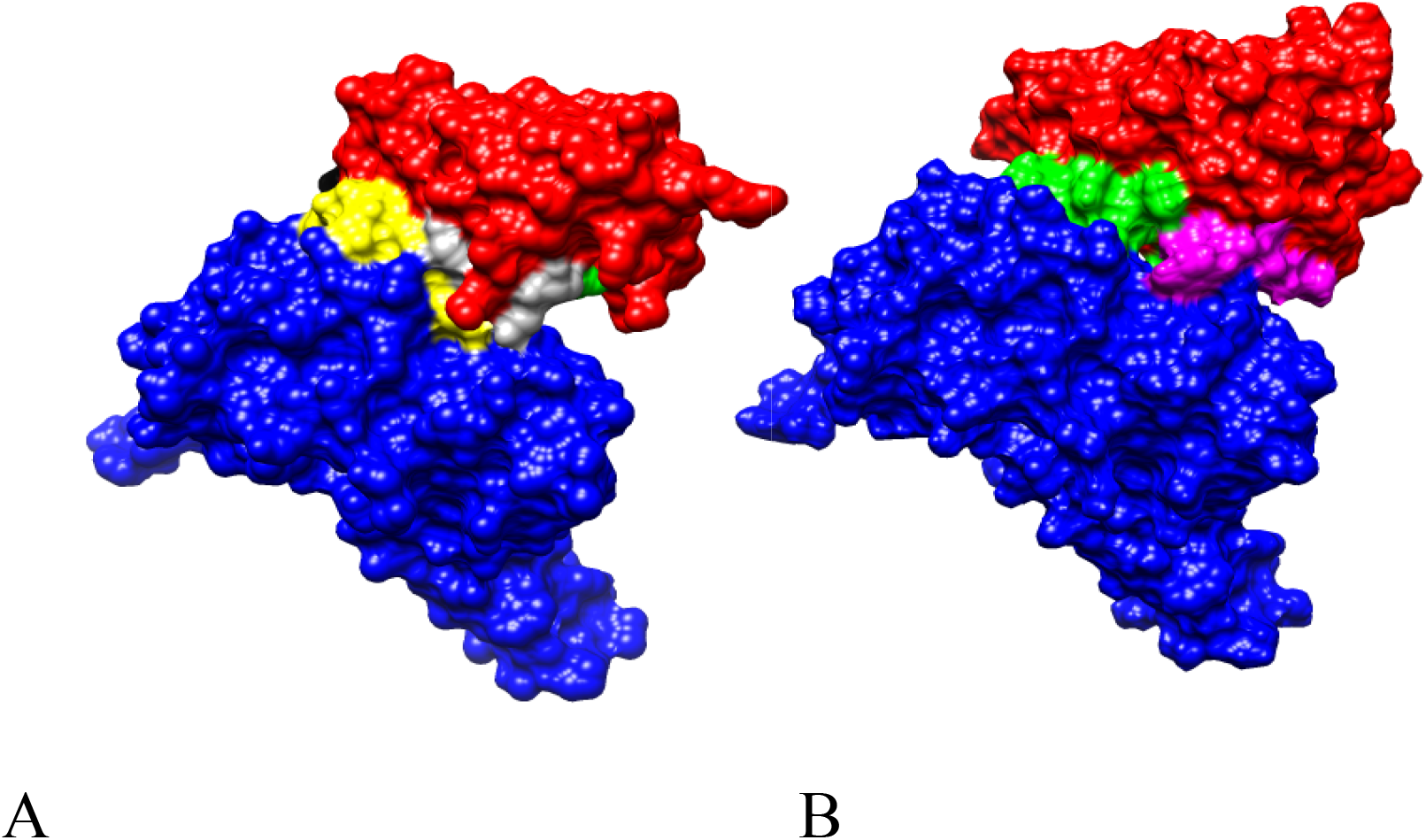
Docked structure of nanobody with RBD A) 6waqM-RBD complex B) 6waqG-RBD complex. **Colour code**: Nanobody-Red, RBD-Blue, CDR1-Black, CDR2-Yellow, FR2-green, FR3-Grey, CDR3-Magenta

## Conclusion

The intricate interrelationship between flexibility, stability and affinity has been subject of intense research, especially in case of antibody. However best of our knowledge this has not been extensively studied in case of nanobodies, although there are reports of stability activity studies (Bekker et al., 2019; Soler et al., 2016; Wang and Duan, 2011) a perspective on all the three simultaneously is the interesting point about this study. Further the results of this study should not be generalized as it is not a benchmarking study and addresses the interrelationship of the three above mentioned variable only in VHH 72. The other caveat in our studies is that we did not consider glycosylation of RBD in our simulation. However studies indicate that of the residues of RBD known to interact with neutralizing antibody 94% were predicted to be antibody accessible (Grant et al., 2020)□. Rigidity in protein will increase favorable enthalpy interactions with decrease in conformational entropy and it has been reported that weakening of HBN far from the mutation site provide enthalpy entropy compensation and counteracting changes in rigidity and flexibility will occur at remote sites to globally balance the rigidity and flexibility in protein structures (Li et al., 2014)□. In antibodies the accumulation of somatic mutation in the CDR and FR regions depends on the maturation pathway. For antibodies, where the germline shows weaker binding to the epitope will accumulate FR mutation to increase the flexibility during early stages and during later stages acquires mutation in the CDR region that contributes to the inceased binding (Ovchinnikov et al., 2018)□. In this study we find the germline binds with lower affinity to the epitope and therefore expected to acquire mutation similar to antibody with germline weaker antigen binding. However a comparison of the framework region in antibody and Nbs shows that FR in nanobody is conserved than antibody (Mitchell and Colwell, 2018). Thus in the absence of partner VL domain and the requirement to remain soluble in isolation provide a strong constraint on which framework sequence changes can be accepted. Thus as per the data in this study mutation in CDR of nbs during the maturation pathway and thereby modulation of the structural flexibility not only is important in functional binding to epitope but may also play a role in fine controling the stability in nanobodies.

## Abbreviations

CDR: Complimentary determining region
Nbs: Nanobodies
FR: Frame work region
RBD: Receptor Binding Domain

